# Landscape-scale grassland structural heterogeneity and disturbance regimes influence swamp deer habitat utilisation across the upper Gangetic plains, India

**DOI:** 10.64898/2026.04.20.719689

**Authors:** Shrutarshi Paul, Sohini Saha, Navendu Page, Bivash Pandav, Samrat Mondol

## Abstract

Large herbivores play a vital role in shaping grassland ecosystems, yet many species face increasing threats from habitat loss, fragmentation, and human disturbance. In northern India, riverine grasslands of the Gangetic floodplains serve as essential habitats for such species, including the swamp deer (*Rucervus duvaucelii duvaucelii*), a habitat specialist that has experienced severe population declines, especially outside protected areas. We evaluated the structural vegetation composition of grasslands and swamp deer habitat use along the Ganges river, covering both protected regions (Jhilmil Jheel Conservation Reserve and Hastinapur Wildlife Sanctuary) and surrounding non-protected grasslands. Combining field-based vegetation surveys with NDVI-based remote sensing enabled the classification of grasslands into four ecologically significant types structured by Saccharum, Phragmites, and Typha species. Disturbance was prevalent across the landscape, with approximately 77% of grassland plots exhibiting signs of human impact. Pure Saccharum grasslands exhibited the highest disturbance levels, followed by mixed patches, while Phragmites*–*Typha patches showed the lowest disturbance. Conversely, Phragmites–Typha patches experienced the lowest levels of disturbance. Data from surveys and two radio-collared swamp deer indicated a consistent preference for mixed vegetation types, particularly Saccharum-dominated and Phragmites–Typha mixes, while avoiding heavily disturbed pure Saccharum patches. Seasonal changes in habitat use reflected context-dependent selection influenced by vegetation structure, hydrology, and disturbance regimes. Our results underscore the importance of conserving diverse grassland mosaics and reducing human pressures, especially in unprotected areas to promote the long-term survival of swamp deer and other grassland herbivores within the Gangetic floodplains.

## Introduction

Large grazing herbivores are vital for maintaining grassland ecosystems (Hobbs 1996, Olff et al. 2002), as their grazing activities play a key role in regulating carbon sequestration and influencing plant species’ richness within these habitats. However, the past century has seen a significant rise in grassland exploitation, fragmentation, and degradation (Kruess and Tscharntke 1994, Fahrig 2003), resulting in the deterioration of about 49% of the world’s grasslands. These changes, along with hunting/poaching and competition with domestic animals for resources, have resulted in a substantial decline in grassland herbivore populations, many of which are now threatened with extinction (Owen-Smith 1989, Lorenzen et al. 2011, Klink et al. 2015). Furthermore, much information on herbivore populations and their habitat status is primarily available from protected areas (Herbert and van der Jeugd 1993, Ahmed and Khan 2008, Bender and Piasecke 2010, Ripple et al. 2016, Dorji et al. 2019, Sinha et al. 2019), while it is lacking from non-protected regions where significant herbivore communities are found (Mallon and Jiang 2009, Karanth et al. 2010, Punjabi and Rao 2017). The future survival and persistence of this species group, their habitats, and associated fauna largely depend on biologically meaningful and accurate information from critical areas. The Indian subcontinent is home to a diverse assemblage of large, grassland-dependent herbivores that face similar declines in population and habitat loss (Karanth et al. 2010). The threatened swamp deer (also known as Barasingha) (*Rucervus duvaucelii*) exemplifies the global conservation challenges faced by habitat-specialist herbivores. The species has experienced over 95% habitat loss in the last century (Karanth et al. 2010, Paul et al. 2020) and currently exists (as three subspecies) within fragmented habitats in northern, central, and eastern India (Sankarshan et al. 1989, Qureshi et al. 2004, Paul et al. 2018, Paul et al. 2020). Populations of the central subspecies (*Rucervus duvaucelii branderi*) and eastern subspecies (*Rucervus duvaucelii ranjitsinhi*) mainly inhabit protected areas and are actively managed (Qureshi et al. 2004, Nath et al. 2023). Conversely, the northern subspecies (*Rucervus duvaucelii duvaucelii*) occurs across a mosaic of protected and unprotected regions in North India (covering the states of Uttarakhand and Uttar Pradesh) and southern Nepal (Khan et al. 2003, Qureshi et al. 2004, Tewari and Rawat 2013b, Paul et al. 2018, Ghimire et al. 2019, Mondol et al. 2019, Paul et al. 2020, Dhami et al. 2023a, Dhami et al. 2023b, Paul et al. 2023a, Paul et al. 2023b), where it faces various conservation challenges. This subspecies is strongly associated with riverine grassland ecosystems along Gangetic floodplains. Riverine grasslands, specifically along the Ganges in India, have experienced approximately 57% habitat loss over the past three decades (Paul et al. 2023a), yet remain critical for the persistence of swamp deer populations outside core protected areas. For large grazers, habitat suitability is strongly influenced not only by plant species composition but also by vegetation structure, which determines forage accessibility, biomass distribution, ease of movement, visibility, and seasonal habitat quality (Denno and Roderick 1991, Fortin et al. 2003, Rech et al. 2025). Structural vegetation patterns are particularly important for species such as swamp deer inhabiting human-dominated areas, as they respond primarily to vegetation structure and biomass distribution, which determine cover, foraging, and birthing efficiency (Tewari and Rawat 2013a, Tewari and Rawat 2013c). An earlier study at a local scale on the swamp deer indicated preferences for mixed vegetation of Saccharum and Typha and avoidance of Phragmites thickets (Tewari and Rawat 2013c). Given the species’ habitat-specialist nature, identifying suitable habitats based on vegetation patterns and structure is crucial for site-specific management and restoration initiatives.

In this paper, we assessed the structural vegetation composition of grassland habitats along the Ganges river (where severe loss was quantified by Paul et al. 2023a) and evaluated its relevance to swamp deer habitat use. The primary focus was on four key grass species: *Saccharum spontaneum*, *Saccharum bengalensis*, *Typha spp*., and *Phragmites spp*., which are essential to the structure of swamp deer habitats and also part of the swamp deer diet. We employed field surveys and GIS-based analyses to identify the unique reflectance signatures of these target species and to assess their spatial coverage, abundance, and disturbance regimes. Additionally, we examined previously available swamp deer movement data (Paul et al. 2023b) to understand swamp deer usage patterns across different grassland types. We believe this detailed information will greatly support future management and restoration efforts in this biodiversity-rich landscape, which is heavily impacted by human activity.

## Materials and methods

### Study area and design

This study was conducted across all the previously identified grassland patches along the river Ganges and its tributaries, covering the states of Uttarakhand and Uttar Pradesh (Paul et al. 2023a), extending from 29°79’99“N, 78°21’71”E to 28°77’34“N, 78°13’57”E. The entire area spans approximately 3173 km², extending from Jhilmil Jheel Conservation Reserve (JJCR) in Uttarakhand to Hastinapur Wildlife Sanctuary (HWLS) in Uttar Pradesh. This area comprises 53% protected areas (HWLS and JJCR, 1677 km²) and 47% unprotected regions (1496 km²). The river Ganges flows through the centre of the study area (around 180 km in length) and is joined by its tributaries Banganga and Solani in Uttar Pradesh. The study encompassed all grasslands located within a maximum distance of eight km from the banks of these three rivers (Supplementary Figure S1). Other land use and land cover (LULC) classes, such as agricultural fields, villages, townships, grassland patches, scrublands, and forests, comprise the majority of this human-dominated landscape. The grasslands are rich in diverse flora (*Saccharum spontaneum*, *Saccharum bengalense*, *Imperata cylindrica*, *Cynodon dactylon*, *Typha angustata*, *Phragmites karka*, *Arundo donax*, etc.) and fauna (swamp deer, hog deer, fishing cat, smooth-coated otter, sarus crane, swamp francolin, etc.) (Paul et al. 2018). This region has been identified as the only primary swamp deer habitat along the Ganges (Paul et al. 2018, Paul et al. 2020), and the fragmented grassland patches are known to be the primary resources available to the subspecies (Khan and Khan 1999, Khan et al. 2003, Qureshi et al. 2004, Tewari and Rawat 2013b, Paul et al. 2018, Mondol et al. 2019, Paul et al. 2020, Paul et al. 2023a, Paul et al. 2023b).

### Assessment of grassland vegetation composition

We employed a combination of field sampling (primary data) and vegetation indices (secondary information) to map (Xie et al. 2008, Kassa et al. 2016) and characterize the main perennial grass species (*Phragmites sp*., *Typha sp*., *Saccharum spontaneum*, and *Saccharum bengalense*) that form the structural framework of swamp deer habitat within this landscape (Tewari and Rawat 2013a, Paul et al. 2020). Initially, we investigated potential differences in reflectance patterns among these major grass species by comparing geo-referenced points in Google Earth Pro imageries (Jamei et al. 2022, Toosi et al. 2022), which indicated potentially varying but unclear patterns of homogeneity or heterogeneity (Supplementary Figure S2). To derive potential species-specific reflectance patterns, we performed a landscape-level vegetation assessment in a phased manner. First, we surveyed all grassland patches between JJCR and Bijnor Barrage of HWLS (upper Gangetic Plains), where 98 training vegetation plots (5×5 m²) were laid out following a stratified random sampling framework (Uribeetxebarria et al. 2019). Stratification and plot number were guided by the degree of vegetation heterogeneity (presence of multiple or single species) observed during field surveys. Within each plot, we counted the individual stems/tufts of *Phragmites sp*., *Typha sp*., *S. spontaneum*, and *S. bengalense*. For homogeneous patches, smaller 1×1 m² plots were laid at each corner of the 5×5 m² plots, and the species were counted and later extrapolated. Subsequently, we classified the plots into pure (≥ 95% coverage of the dominant species) or dominated mix (at least 25% coverage of the dominant species) vegetation classes (Wikum and Shanholtzer 1978) as follows:

1) Pure *Typha* sp.
2) Pure *Phragmites* sp.
3) Pure *Saccharum* (consists of pure *S. bengalense* and/or *S. spontaneum*)
4) *Typha* Dominated Mix (dominated by *Typha* sp. with the presence of three other species)
5) *Phragmites* Dominated Mix (dominated by *Phragmites* sp., with the presence of the other three species)
6) *Saccharum* Dominated Mix (dominated by *S. bengalense* and/or *S. spontaneum* with presence of *Phragmites* sp. */ Typha* sp.)

Further, we used the field-collected vegetation data to generate a Normalized Difference Vegetation Index (NDVI) based map (Bannari et al. 1995, Eastman et al. 2013, Elgammal et al. 2014). For this purpose, we downloaded Landsat images (Landsat 8-USGS, 2018) with 30 m resolution (Details in Supplementary Table S1) and chose visible red (band 4) and near-infrared (band 5) wavelengths to calculate NDVI using the formula-

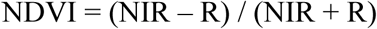

where R and NIR imply surface reflectance averaged over red (λ – 0.6 μm) and near-infrared (NIR) (λ – 0.8 μm) regions of the spectrum, respectively (Jiang et al. 2006, Elgammal et al. 2014, Aswatha et al. 2018). Once the raw NDVI data were generated, we calculated the class-specific NDVI ranges for each vegetation plot, followed by extrapolation to the grassland areas only (Paul et al. 2023b) using the Reclassify Tool in ArcGIS 10.2.2 (Beckham and Atkinson 2017).

Finally, we conducted an extended vegetation sampling from south of Bijnor Barrage to the southern boundary of HWLS (111 plots) to estimate the accuracy of NDVI classification. The accuracy was calculated using the following formula:

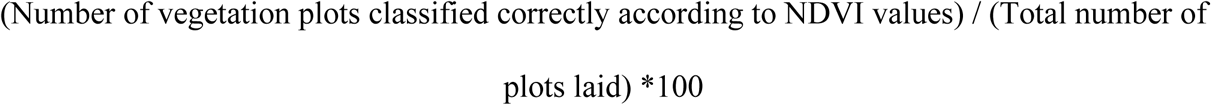

We also gathered information on human disturbances (such as lopping, cutting, and grazing) from each plot (both training and test plots) and classified them according to disturbance levels (low, moderate, and high). We assessed whether disturbance levels differed among vegetation types using an ordinal logistic regression (polr function in MASS package in R) (Ripley and Venables 2023) with disturbance as the ordered response variable (Low < Moderate < High) and vegetation type as a predictor. The vegetation type exhibiting the lowest overall disturbance level was designated as the reference category. Coefficients were exponentiated to obtain odds ratios (ORs) with 95% confidence intervals. OR > 1 indicates higher odds of disturbance relative to the least disturbed vegetation type.

We used ArcGIS 10.2.2 to estimate the area coverage of each vegetation type (based on NDVI values) across the study area. To evaluate differences in vegetation composition among the sampled plots, we performed Non-metric Multi-Dimensional Scaling (NMDS) using Scatterplot3d (Oksanen 2015) and the Vegan package in R, (https://CRAN.R-project.org/package=vegan) and tested the significance of differences among vegetation types with a bootstrapping PERMANOVA test. Additionally, we investigated the relative abundance of each major grass species (based on field data) within each vegetation type. The Kruskal-Wallis test was used to assess significant differences among these species (Kruskal 1964, Reinecke et al. 2014).

### Swamp deer vegetation preferences

We employed a combination of radio-tracking and field surveys to understand swamp deer’s vegetation-type preferences. We fitted GPS-enabled radio collars on two healthy female swamp deer at JJCR in May-June 2018 and tracked them for 11 months (Female 1) and 14 months (Female 2), respectively. The collaring was performed using a drive-net approach with all necessary precautions during field operations (see Paul et al. 2023, for details). The collars were set to provide information on latitude, longitude, time, and temperature every two hours. To understand grassland type preferences, we combined the location points and home range data of the collared females (see Paul et al. 2023) with earlier information on swamp deer presence (based on antlers and genetic data) from the entire landscape (Paul et al. 2018, Paul et al. 2020, Paul et al. 2023a, Paul et al. 2023b). We calculated the proportions of different vegetation types within the 50% and 95% BBMM home ranges (representing the core area of use and standard home range size). We assessed possible seasonal (summer, monsoon, and winter) variations in vegetation preferences using Ivlev’s electivity index (Ivlev 1961). The home range analysis was conducted at a resolution of 30 metres using the BBMM package (Nielson and McDonald 2015) in R software (Horne et al. 2007).

We hypothesized the following:

a) Grassland regions dominated by *Saccharum* are likely more widespread in this area, as its presence is expected to be greater along rivers. Factors such as barrages, flooding events, and river dynamics may have aided the colonisation of barren lands along the Ganges by *Saccharum* (Paul et al. 2023).
b) Anthropogenic disturbance is expected to vary across vegetation types, with structurally open or more accessible grasslands (e.g., *Saccharum* stands) experiencing higher levels of grazing, cutting, and lopping than denser *Phragmites–Typha* stands.
c) Swamp deer may not show specific preferences for any grassland throughout the year, due to the highly fragmented and degraded habitats present in this landscape, especially when compared to more protected areas.

## Results

### Species composition of the grasslands

Based on the classification criteria described before (see methods), we assigned the training vegetation plots (between JJCR and Bijnor Barrage of HWLS, n=98) into following: Pure *Typha*-five plots; Pure *Phragmites*-29 plots; Pure *Saccharum*-19 plots; Typha Dominated Mix-one plot, *Phragmites* Dominated Mix-23 plots; and *Saccharum* Dominated Mix-21 plots, respectively. We generated associated NDVI value ranges from these assigned plots, and during the generation of NDVI-based vegetation classifications, we ascertained that Phragmites and Typha could not be separated due to their close association in the field and similar NDVI profiles. Therefore, we reclassified the training plots based on a combination of species-specific data and associated NDVI values (from already surveyed training plots) into the following four categories with respective NDVI ranges: Pure *Saccharum* (PS; NDVI:0.12-0.315), *Saccharum* Dominated Mix (SDM; NDVI:0.316-0.39), Phragmites-Typha Dominated Mix (PTDM; NDVI:0.40-0.44) and Pure Phragmites-Typha (PPT; NDVI:0.45-0.56) (Fig. 1b). We obtained 86.4% accuracy in correctly classifying the sampled plots (n=111) between south of Bijnor Barrage and the southern boundary of HWLS (Supplementary Table S2). Finally, we had 194 plots and removed the incorrectly classified plots (n= 15) from further downstream analysis. Pure *Saccharum* was the dominant vegetation type covering 35% of the total grassland area (144.04 km²), followed by Pure Phragmites-Typha (27%), *Saccharum* Dominated Mix (23%) and Phragmites-Typha Dominated Mix (15%). Similar vegetation-type trends were observed when comparing protected and non-protected areas within this landscape (Table 1).

**Fig. 1.**
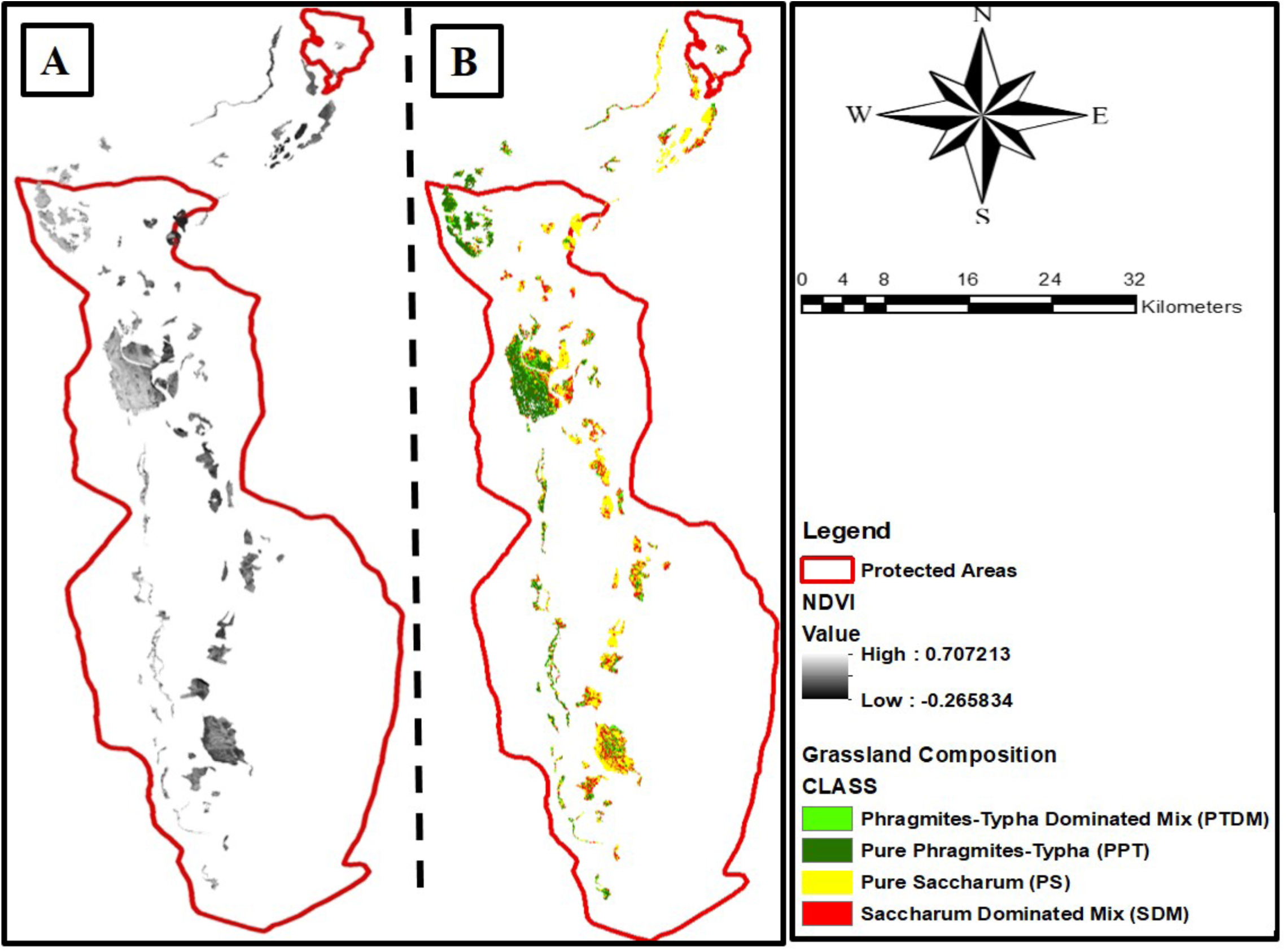
Structural vegetation classification within grasslands in the upper Gangetic plains: A) Raw NDVI value ranges between-0.2 and +0.7, where very low values indicate water bodies, barren areas, and sands; moderate values specify shrub and grassland. B) NDVI-based reclassification of grassland vegetation types into four categories: Pure Saccharum (PS; 0.12–0.315), Saccharum Dominated Mix (SDM; 0.316–0.39), Phragmites–Typha Dominated Mix (PTDM; 0.40–0.44), and Pure Phragmites–Typha (PPT; 0.45–0.56), based on species composition and associated NDVI value ranges

**Table 1.** Area (km²) of grassland within protected and non-protected areas, and their percentage contribution.

| **Vegetation Class** | **Grassland area in Total landscape (Km<sup>2</sup>)** | **Grassland area in Protected Area (Km<sup>2</sup>)** | **Grassland area in Non-Protected Area (Km<sup>2</sup>)** |
| --- | --- | --- | --- |
| Pure Saccharum (PS) | 50.73 (35%) | 39.31 (32%) | 11.42 (53%) |
| Saccharum Dominated Mix (SDM) | 32.76 (23%) | 27.95 (23%) | 4.81 (22%) |
| Phragmites-Typha Dominated Mix (PTDM) | 21.77 (15%) | 19.43 (16%) | 2.34 (11%) |
| Pure Phragmites-Typha (PPT) | 38.76 (27%) | 35.74 (29%) | 3.02 (14%) |
|  | 144.02 | 122.43 | 21.59 |

Out of 194 (98 training+96 correctly classified from test plots) vegetation plots, 149 (77%) plots were found to retain signatures of high disturbances, whereas 22% (n=42) and 1% (n=3) showed moderate and low disturbances, respectively. Out of the four vegetation classes, plots classified under high disturbance were most frequently associated with pure *Saccharum* (n = 42, representing 93% of plots in the high-disturbance category), whereas the Pure *Phragmites–Typha* class contributed the lowest proportion of highly disturbed plots (n = 29, 59%) (Supplementary Figure S3). When comparing disturbance regimes across plots, we found PS and SDM had much higher odds of being in a higher disturbance category than PPT (the least disturbed category), with odds ratios of ∼10 and 5.8, respectively (p < 0.01). In contrast, PTDM showed slightly higher odds (OR∼1.2) that were not significant (p>0.01) (Details in Table 2).

**Table 2.** Ordinal logistic regression results from a proportional odds model assessing the impact of vegetation type on disturbance levels, with Pure Phragmites–Typha (PPT) as the reference category. Odds ratios (OR) indicate the change in odds of higher disturbance compared to the reference vegetation type.

| **Vegetation Type** | **Coefficient ($\beta$)** | **Std. Error** | **t-value** | **p-value** | **Odds Ratio (OR = $\exp(\beta)$)** | **95% CI (OR)** | **Interpretation** |
| --- | --- | --- | --- | --- | --- | --- | --- |
| PPT (Pure Phragmites–Typha) | – | – | – | – | 1 | – | Reference (least disturbed) |
| PTDM (Phragmites–Typha Dominated Mix) | 0.154 | 0.437 | 0.353 | 0.724 | 1.17 | 0.50 – 2.75 | More disturbed than PT, not significant |
| PS (Saccharum) | 2.325 | 0.665 | 3.496 | 0.00047 | 10.23 | 2.78 – 37.66 | Much more disturbed than PT, significant |
| SDM (Saccharum-Dominated Mix) | 1.75 | 0.496 | 3.528 | 0.00042 | 5.76 | 2.18 – 15.22 | More disturbed than PT, significant |

NMDS ordination revealed three separate vegetation clusters (Stress value 0.13), supporting differences in species composition among the four vegetation classes identified in this landscape (PERMANOVA, p < 0.01) (Fig. 2). The Saccharum and Saccharum-Dominated Mix vegetation classes overlapped with each other. Within the Saccharum class, *S. spontaneum* was the most abundant among the four species. Saccharum Dominated Mix and Phragmites Dominated Mix classes showed abundance of respective species within each plot type, along with other species. The most abundant species in the Phragmites-Typha class was *Phragmites* sp., followed by *Typha* sp. and *Saccharum* spp. (Fig. 3). Comparison of the mean relative abundance of each species across the four vegetation types also revealed significant differences (p < 0.01, Kruskal-Wallis).

**Fig. 2.**
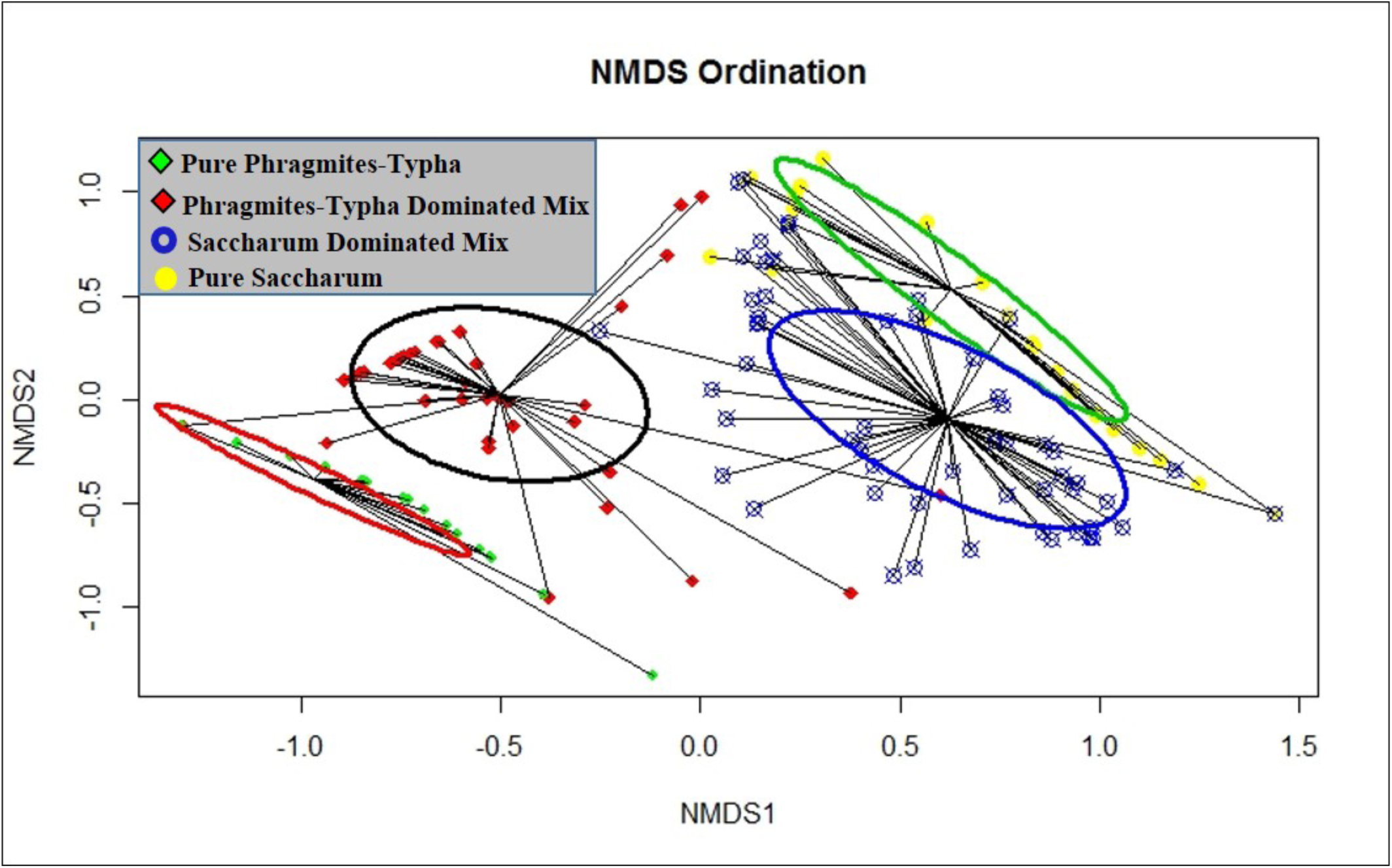
Non-metric Multidimensional Scaling analysis displaying grassland vegetation classes in two dimensions, where green, red, blue, and yellow represent Pure Phragmites-Typha (PPT), Phragmites-Typha Dominated mix (PTDM), Saccharum Dominated Mix (SDM), and Pure Saccharum (PS) respectively

**Fig. 3.**
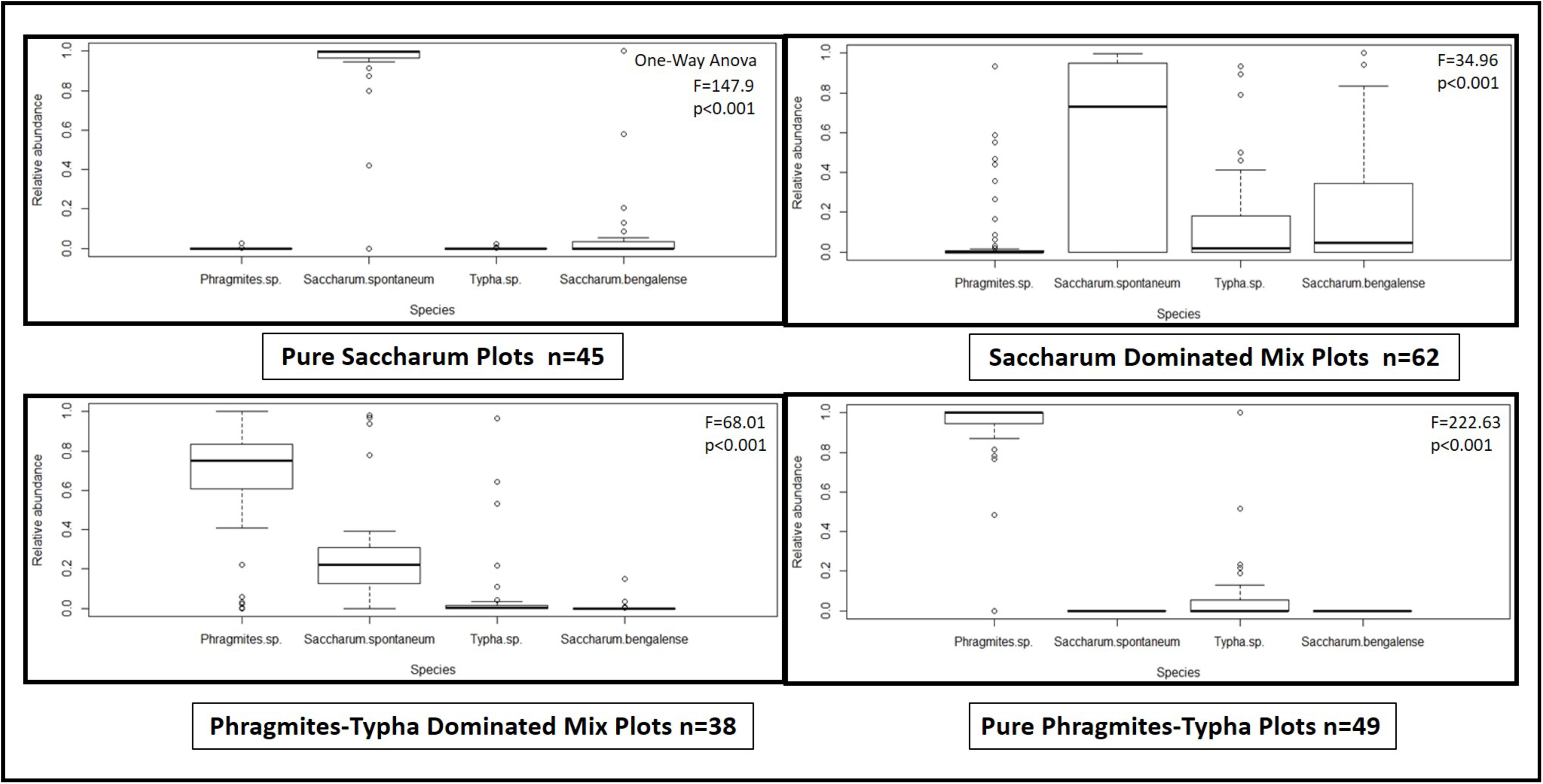
Relative abundance of each species (*Phragmites* sp., *Saccharum spontaneum*, *Saccharum bengalense*, and *Typha* sp.) within the four vegetation classes: A) Pure Saccharum (PS), B) Saccharum Dominated Mix (SDM), C) Phragmites-Typha Dominated Mix (PTDM), and D) Pure Phragmites-Typha (PPT)

### Grassland vegetation preference of the swamp deer

We obtained a total of 9,159 GPS locations from both radio-collared individuals, and most of them (∼81%, n = 7419 location points) were found within the grasslands. Upon close inspection, we identified that ∼57% (n=4229) of the grassland GPS locations were from Saccharum Dominated Mix vegetation type, followed by Pure Saccharum (∼25%, n=1855), Phragmites-Typha Dominated Mix (∼13%, n=964) and Pure Phragmites-Typha (∼5%, n=371). In addition, we also used a total of 466 GPS location details from males (n=395), females (n=62) and unidentified (n=9) (collected during earlier surveys, Paul et al. 2023b) across this landscape. Among the fecal samples collected during survey, approximately 30% (n = 144) originated from SDM vegetation, followed by PPT (∼29%, n = 138), PTDM (∼27%, n = 128), and PS (∼12%, n = 56). Among samples attributed to males, the highest proportion came from PPT vegetation (31%, n = 125), followed by PTDM (∼29%, n = 117), SDM (∼28%, n = 113), and PS (∼10%, n = 40). For females, most fecal samples were collected from SDM vegetation (45%, n = 28), followed by PS (∼22%, n = 14), PPT (∼17%, n = 11), and PTDM (∼14%, n = 9).

The average home ranges for the two individuals were 10.27 km^2^ and 1.02 km^2^ for the 95% (and 50% (core home range) BBMM, respectively. The habitat preferences of both individuals showed that the collared females preferred SDM vegetation but avoided PS vegetation (Fig. 4, Table 3). At the individual level, Female 1 preferred SDM vegetation, along with some selection of PTDM (95% and 50% BBMM scales) and PPT (50% BBMM scale), whereas Female 2 preferred only SDM across all scales. At the seasonal level, results indicate that both collared individuals preferred PTDM vegetation in summer and SDM in winter but avoided PS in both seasons. However, during the monsoon, both individuals showed avoidance towards PPT and PTDM, while mostly preferring PS and SDM (Details in Table 3).

**Fig. 4.**
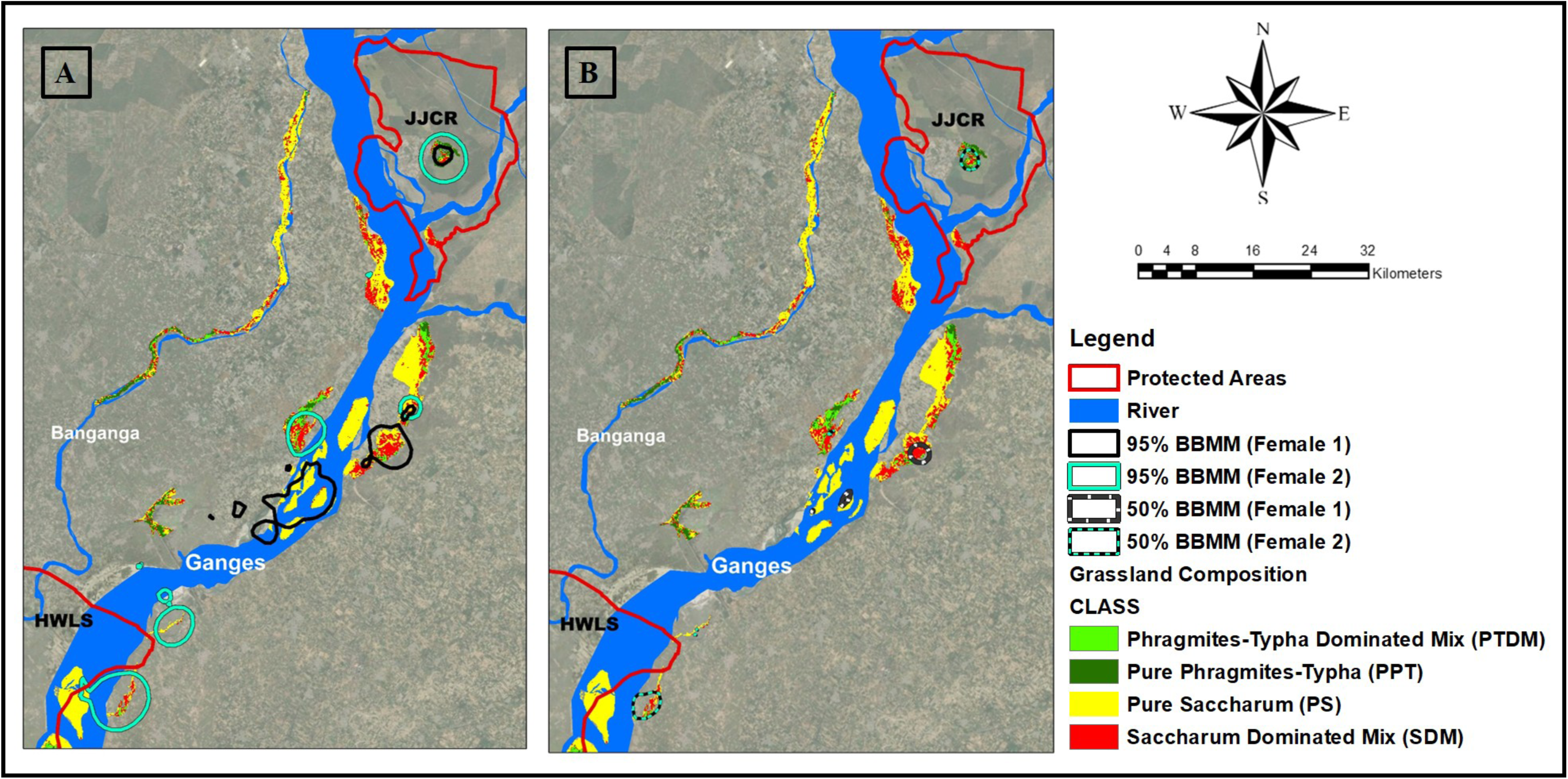
Habitat preference patterns of collared females (Female 1 & 2) across vegetation classes at 95% home range and core home range (50%) levels. Both females showed strong selection for SDM vegetation and avoidance of PS vegetation. At the individual level, Female 1 additionally selected PTDM (at both 95% and 50% BBMM scales) and PPT (at 50% BBMM scale), whereas Female 2 exclusively preferred SDM across all scales

**Table 3.** Seasonal habitat preference (Ivlev’s index) of female collared individuals across vegetation classes within 95% home range and 50% BBMM home ranges.

|  | **Whole year** |  |
| --- | --- | --- |
|  | **Female 1** | **Female 2** |
| PS (50% BBMM/ 95% BBMM) | (0.13/0.28) | (0.11/0.19) |
| SDM (50% BBMM/ 95% BBMM) | (0.68/0.56) | (0.63/0.58) |
| PTDM (50% BBMM/ 95% BBMM) | (0.13/0.11) | (0.16/0.15) |
| PPT (50% BBMM/ 95% BBMM) | (0.04/0.03) | (0.08/0.06) |
|  | **Summer** |  |
|  | **Female 1** | **Female 2** |
| PS (50% BBMM/ 95% BBMM) | (0.03/0.16) | (0.12/0.12) |
| SDM (50% BBMM/ 95% BBMM) | (0.79/0.67) | (0.35/0.36) |
| PTDM (50% BBMM/ 95% BBMM) | (0.12/0.10) | (0.29/0.30) |
| PPT (50% BBMM/ 95% BBMM) | (0.04/0.04) | (0.21/0.20) |
|  | **Winter** |  |
|  | **Female 1** | **Female 2** |
| PS (50% BBMM/ 95% BBMM) | (0/0.007) | (0.20/0.23) |
| SDM (50% BBMM/ 95% BBMM) | (0.74/0.74) | (0.77/0.71) |
| PTDM (50% BBMM/ 95% BBMM) | (0.17/0.19) | (0.02/0.04) |
| PPT (50% BBMM/ 95% BBMM) | (0.07/0.05) | (0/0) |
|  | **Monsoon** |  |
|  | **Female 1** | **Female 2** |
| PS (50% BBMM/ 95% BBMM) | (0.70/0.72) | (0.12/0.15) |
| SDM (50% BBMM/ 95% BBMM) | (0.25/0.23) | (0.66/0.66) |
| PTDM (50% BBMM/ 95% BBMM) | (0.03/0.03) | (0.20/0.18) |
| PPT (50% BBMM/ 95% BBMM) | (0/0) | (0/0) |

## Discussion

Our assessment of grassland vegetation and swamp deer along the Ganges between JJCR and HWLS provides insights into the species-vegetation dynamics and the effects of human disturbance across protected and unprotected areas. The dominance of *Saccharum* grasslands and exposure to high human disturbance highlight ecological challenges for grassland fauna. Results also show swamp deer use more diverse mixed patches, underscoring the importance of grassland diversity for their survival, especially outside protected zones. Vegetation mapping, NDVI classifications and field validations revealed that about 35% of the grasslands are dominated by pure Saccharum, followed by mixed types and pure Phragmites-Typha. Such patterns reconfirmed earlier suggestions that *Saccharum* spp. are successful colonisers in empty riverine areas disturbed by flooding, river changes, or human activities such as barrages (Dinerstein 2003, Paul et al. 2023b). However, 77% of grasslands are disturbed, with pure *Saccharum* patches experiencing the most human impact, likely due to their accessibility and use for fodder, thatch, or land conversion (Tarhouni et al. 2010, Pandey et al. 2015, Hassan et al. 2024) as hypothesized. Conversely, Phragmites-Typha patches experienced the least disturbance, possibly due to their dense growth, year-long inundations and lower palatability compared to forage grasses (Kiviat 2013, Durant et al. 2020). Such disturbance regimes are concerning, as ongoing human pressure can reduce habitat complexity, alter species composition, and weaken grassland resilience (Kruess and Tscharntke 1994, MacDougall et al. 2013, Smith et al. 2022). Analyses of combined field botanical data with NDVI spectral information have enhanced grassland mapping efforts in dynamic riverine landscapes and refined vegetation classifications specific to swamp deer. We initially classified six vegetation types (based on available information) and finally consolidated them into four major, swamp deer-specific, ecologically meaningful vegetation categories (due to spectral overlap between Phragmites and Typha): Pure Saccharum, Saccharum Dominated Mix, Phragmites-Typha Dominated Mix, and Pure Phragmites-Typha. This situation also underscores a common limitation in remote sensing, where key species are spectrally indistinguishable despite ecological differences (Xie et al. 2008, Pesaresi et al. 2020). In this study, NDVI class thresholds derived from rigorously sampled field plots enabled us to achieve an overall accuracy of 86.4%, confirming the reliability of our classification in representing the dominant grassland types relevant to swamp deer in this heterogeneous landscape. Furthermore, the NMDS results provided robust multivariate evidence supporting the vegetation classification based on field and remote sensing, increasing confidence that these classes represent ecologically meaningful habitat units. The intensive radio-telemetry locations and field data provided the first quantitative account of grassland vegetation preferences of the northern swamp deer in the Gangetic floodplains. Contrary to our initial hypothesis of non-selectivity due to habitat fragmentation, swamp deer favoured mixed grassland habitats, which potentially offer diverse structures, better protection, and foraging options (Fuhlendorf and Engle 2001, Owen-Smith 2002, Tewari and Rawat 2013a). The slightly greater use of Saccharum-dominated mix compared to Phragmites-dominated mix may reflect marginal differences in forage quality and structural openness, with Saccharum stands likely offering more accessible and palatable resources rather than indicating strong habitat specialisation (Owen-Smith 2002). Avoidance of Pure Saccharum habitats by two collared individuals may be due to heavy disturbance in these habitats (Lv et al. 2024). The use of mixed vegetation types may reflect the swamp deer’s adaptive behaviour in balancing resource availability and risk. We believe that in landscapes with limited or degraded prime habitats, swamp deer may prefer mosaics that provide both sufficient forage (*Saccharum* and *Typha*) and cover (*Phragmites*, *Typha),* similar to patterns observed in other large grassland ungulates facing disturbance regimes (Fuhlendorf and Engle 2001, Ripple et al. 2015). The ecological importance of these complex habitat mosaics for swamp deer can be better understood by examining their seasonal use patterns. A detailed analysis of habitat use revealed seasonal shifts in grassland preferences, with both collared female swamp deer favouring mixed habitat patches (Saccharum-dominated and Phragmites-Typha mix) over pure Saccharum areas during summer and winter. However, during the monsoon, their habitat choices change to favour pure Saccharum types, probably because waterlogging in Typha/Phragmites zones (shallower regions) makes Saccharum habitats more suitable for movement and birthing purposes (Paul et al. 2023b). Moreover, less disturbance during the monsoons allows swamp deer easier access to sites dominated by Saccharum spp. *Saccharum spontaneum* is known to thrive in moist, fertile soils but prefers well-drained or seasonally wet sites over permanently waterlogged ones (Sankaran et al. 2004, Rawat and Adhikari 2015). Earlier studies have indicated that northern swamp deer favour Typha-dominated marshes in relatively undisturbed environments and exhibit seasonal migration patterns for birthing purposes (Tewari and Rawat 2013c, Paul et al. 2023b). The findings from this study suggest a more complex, context-dependent habitat selection pattern driven by availability, seasonal differences and disturbance regimes within this human-dominated landscape. As these patterns are based on female swamp deer, future studies should focus on male habitat preferences in this landscape to gain in-depth information on sex-specific habitat preferences, if any. Although no males were GPS-collared in the present study, opportunistic evidence from fecal samples (genetically identified as males) and shed antlers (Paul et al. 2023a, Paul et al. 2023b) indicated that males also used mixed wetland–grassland patches. In addition, camera trap records from these grasslands frequently detected males in association with female groups within relatively less fragmented patches. Together, these observations suggest that males may also rely on these habitat types, but targeted telemetry or systematic sampling will be required to confirm fine-scale sex-specific selection patterns.

This study provides valuable management insights into the habitat’s structural composition and the use of swamp deer along the Gangetic plains. The abundance of Saccharum, especially *S. spontaneum*, along altered riverbanks suggests certain grass species take advantage of disturbed, open, or flooded sites (Paul et al. 2020). However, without active restoration, the sustainability of such opportunistic regeneration is uncertain, especially given a 57% loss of Gangetic grasslands over the past 30 years (Paul et al. 2023a). Recent records of anthropogenic disturbances in Saccharum-dominated habitats (Paul et al. 2023a) suggest the need for a landscape-level management plan to address resource extraction, prevent conversion, and restore degraded patches, particularly in unprotected regions with swamp deer populations. Although our data suggest relatively lower disturbances in Pure Phragmites-Typha patches, these areas have been fragmented over the past 30 years, especially along the Solani River, a Ganges tributary (Paul et al. 2023a). River dynamics, both natural and influenced by human infrastructure such as dams and barrages, continue to create new grassland patches but may also unpredictably remove or isolate key habitats. Therefore, an adaptive, landscape-scale management strategy involving year-round monitoring of grassland changes to identify and respond to rapid shifts will be vital for long-term habitat preservation. Importantly, swamp deer in this landscape showed higher use of mixed-vegetation patches than of homogeneous stands, indicating that structurally heterogeneous grassland–wetland mosaics likely provide a combination of forage diversity, cover, and microhabitat conditions (Cromsigt et al. 2009, László et al. 2018). Maintaining such heterogeneity should therefore be a management priority. This can be supported through rotational or patch-based disturbance regimes (e.g., controlled seasonal cutting or burning where appropriate), prevention of single-species dominance, regulated domestic cattle grazing, etc. Additionally, improving connectivity between remaining grassland patches in both protected and unprotected areas is crucial for maintaining ecological integrity and securing the long-term survival of swamp deer along the Jhilmil-Hastinapur corridor. Creating riparian buffer zones and community reserves can act as wildlife corridors, stabilise riverbanks, and support habitat mosaics needed for seasonal movement, while also considering the region’s socio-economic realities. Many households in the Jhilmil–Hastinapur corridor depend on seasonal grazing, fodder collection, and riverine resources for their livelihoods. Conservation planning should therefore incorporate participatory management frameworks, negotiated grazing schedules, and the designation of shared-use zones to balance ecological integrity with livelihood security. Incentive-based mechanisms, including compensation schemes, alternative fodder provisioning, and support for sustainable income-generating activities (e.g., community-based eco-tourism, agroforestry), may reduce pressure on core swamp deer habitats. Multi-stakeholder collaboration that integrates awareness programs and livelihood-support strategies can help mitigate anthropogenic pressures on prime swamp deer habitats.

Although the study provides pioneering insights into landscape-scale grassland habitats in the upper Gangetic plains, it is essential to acknowledge certain limitations associated with this research. For example, despite a detailed understanding of vegetation structure and habitat utilisation, the study did not examine the dietary composition of northern swamp deer due to logistical challenges. Future research should incorporate assessments of diet composition and nutritional quality to better understand swamp deer habitat suitability and foraging behaviour. Additionally, this study is based on data collected over one year from two females. Although the NDVI signatures of grassland types showed consistent and reliable patterns throughout the year (Supplementary Figure S4), multi-seasonal sampling (along with telemetry data) is necessary to significantly enhance understanding of the dynamic changes in vegetation composition, structure, and availability across the Gangetic floodplain. Furthermore, we examined four main grass species in the context of swamp deer use; however, other species, such as *Cynodon* sp. and *Imperata* sp., also contribute to habitat diversity (Tewari and Rawat 2013c). Future studies incorporating a wider range of vegetation components would offer a more comprehensive characterisation of the habitat, ultimately benefiting the overall grassland fauna in this landscape.

In summary, this study highlights the vital role of structurally and compositionally diverse grassland mosaics in supporting the persistence of swamp deer in the rapidly changing Gangetic landscapes. We believe the challenges faced by swamp deer are similar to those encountered by many grassland herbivores worldwide, making our findings relevant for human-dominated landscapes with fragmented habitat patches. Combining ecological data-driven monitoring with policy measures that account for socio-economic conditions offers the best chance of sustaining large herbivore populations in unprotected areas globally.

## Acknowledgement

We acknowledge the forest departments of Uttarakhand and Uttar Pradesh and the Ministry of Environment, Forest and Climate Change for granting research permits. We are grateful to the Divisional Forest Officer of the Haridwar Forest Division and local community members for their assistance during fieldwork. We sincerely acknowledge Dr. Parag Nigam and Dr. Bilal Habib for their invaluable contributions to the collaring operation. Field and logistical support from Vinod Thakur, Tista, Suvankar, Dr Sarbesh, Shiv, Shrushti, Amir, Kaushal, Sultan, Nimisha, Rakesh, Debanjan, Prajak, Chitrapal, Imam, Ranju, Bhura, Annu, Juri, Inam, Ammi, and the master’s students of the Wildlife Institute of India (XVI batch) is sincerely appreciated. We also acknowledge technical assistance in GIS work from Dr Gautam, Zeeshan, Aishwariya, and Shaheer. Institutional support from the Director, Dean, and Research Coordinator of the Wildlife Institute of India is gratefully recognized.

## Author Contributions

SP: Conceptualization, Methodology, Data curation, Investigation, Formal analysis, Visualization, Writing – Original draft, Writing – Review and editing. SS: Methodology, Data curation, Formal analysis, Visualisation, Writing – Review and editing. NP: Conceptualization, Writing – Review and editing, Supervision. BP: Conceptualization, Writing – Review and editing, Funding acquisition, Resources, Supervision, Project administration. SM: Conceptualization, Writing – Original draft, Writing – Review and editing, Funding acquisition, Resources, Supervision, Project administration, Investigation.

## Competing Interests

The authors declare no competing interests.

## Data Availability

Additional data from this study are available from the corresponding authors on request.

## Supplementary material legends

**Supplementary Figure S1.**
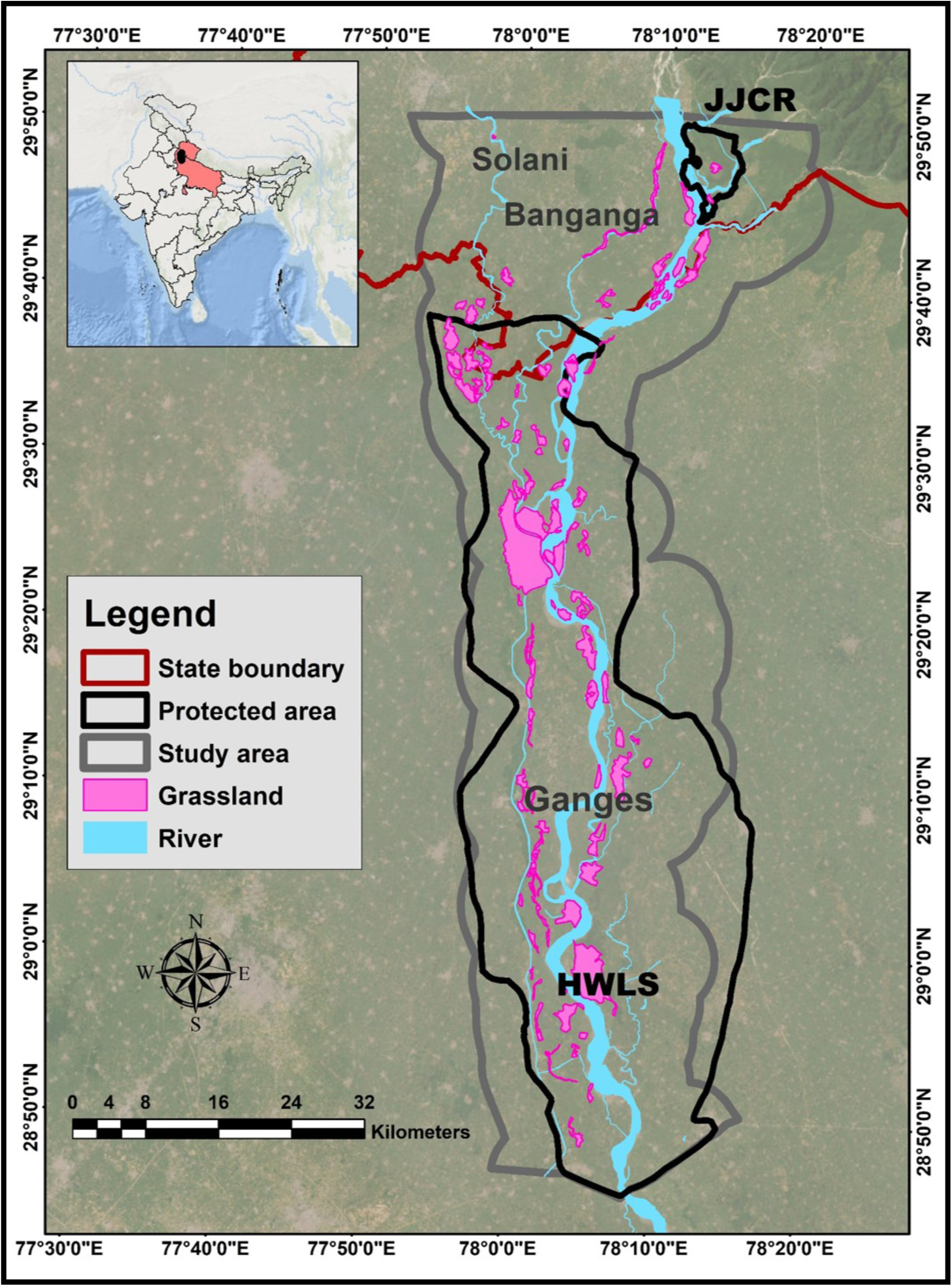
Map of the study area showing the grassland patches. The study area consists of two protected areas (JJCR and HWLS) and other non-protected regions.

**Supplementary Figure S2.**
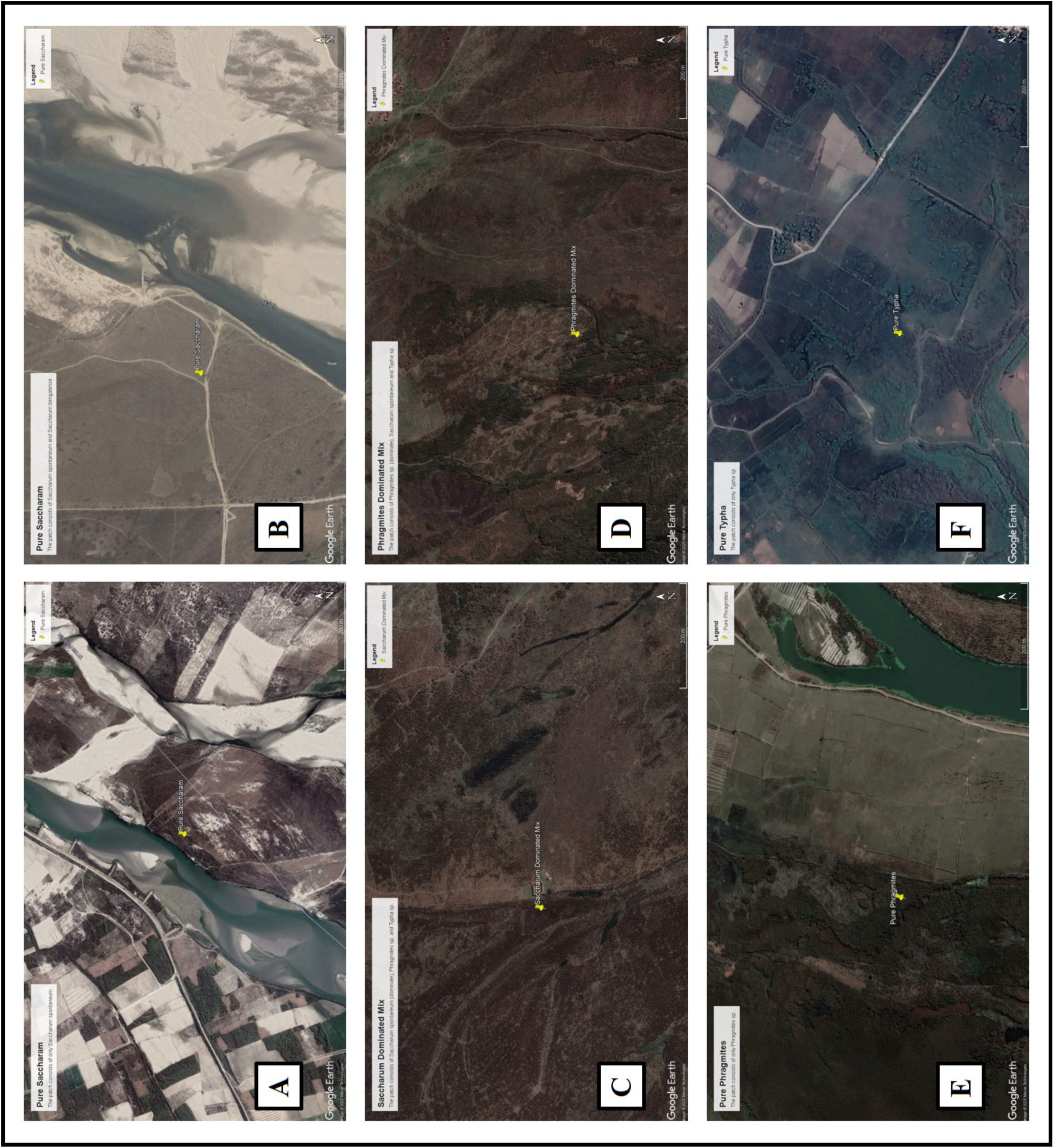
Comparison of reflectance patterns among major grass species using geo-referenced points in Google Earth Pro imagery, indicating potential variation in homogeneity and heterogeneity across vegetation types.

**Supplementary Figure S3.**
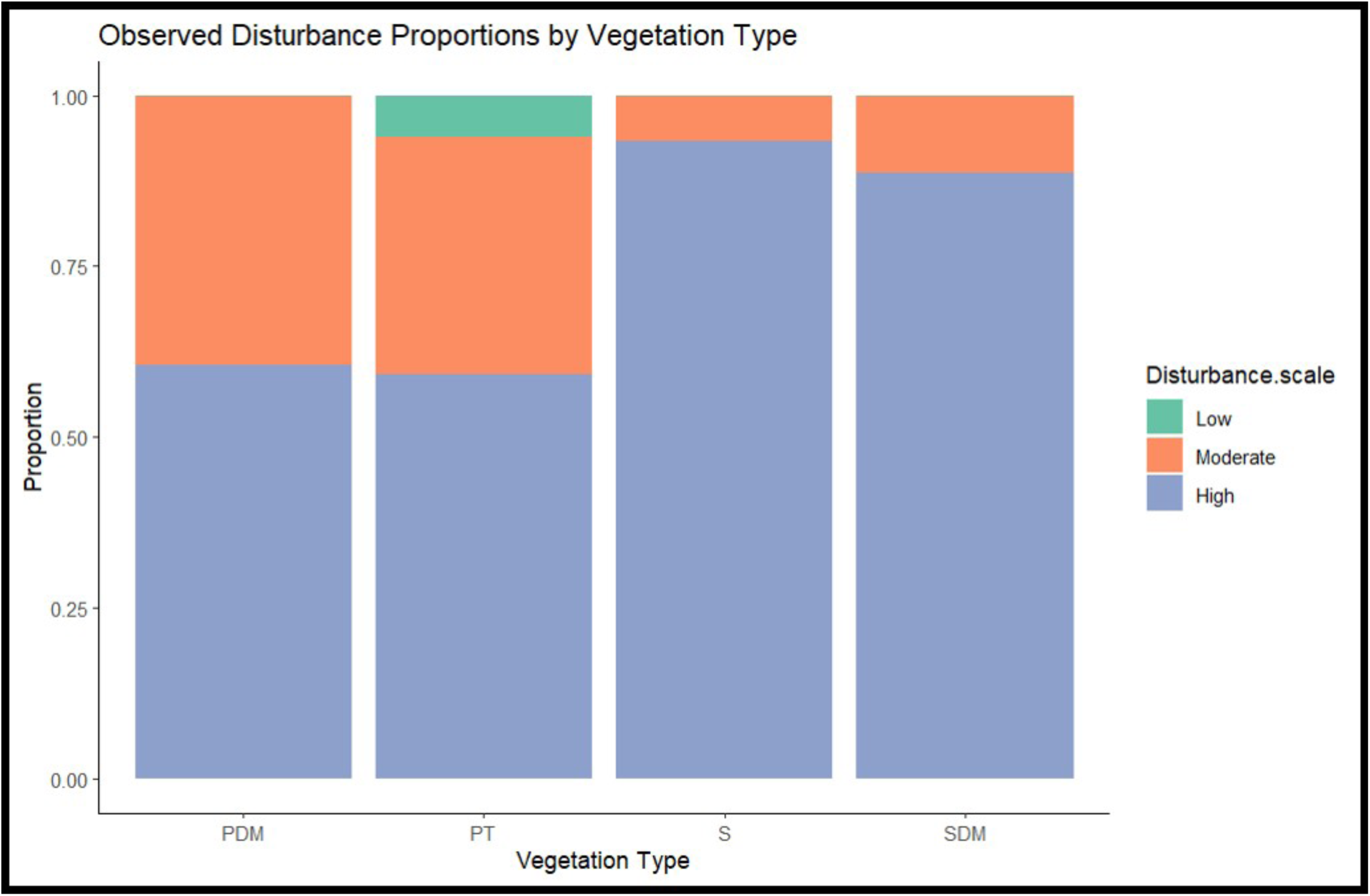
Differences in disturbance regimes among vegetation classes.

**Supplementary Figure S4.**
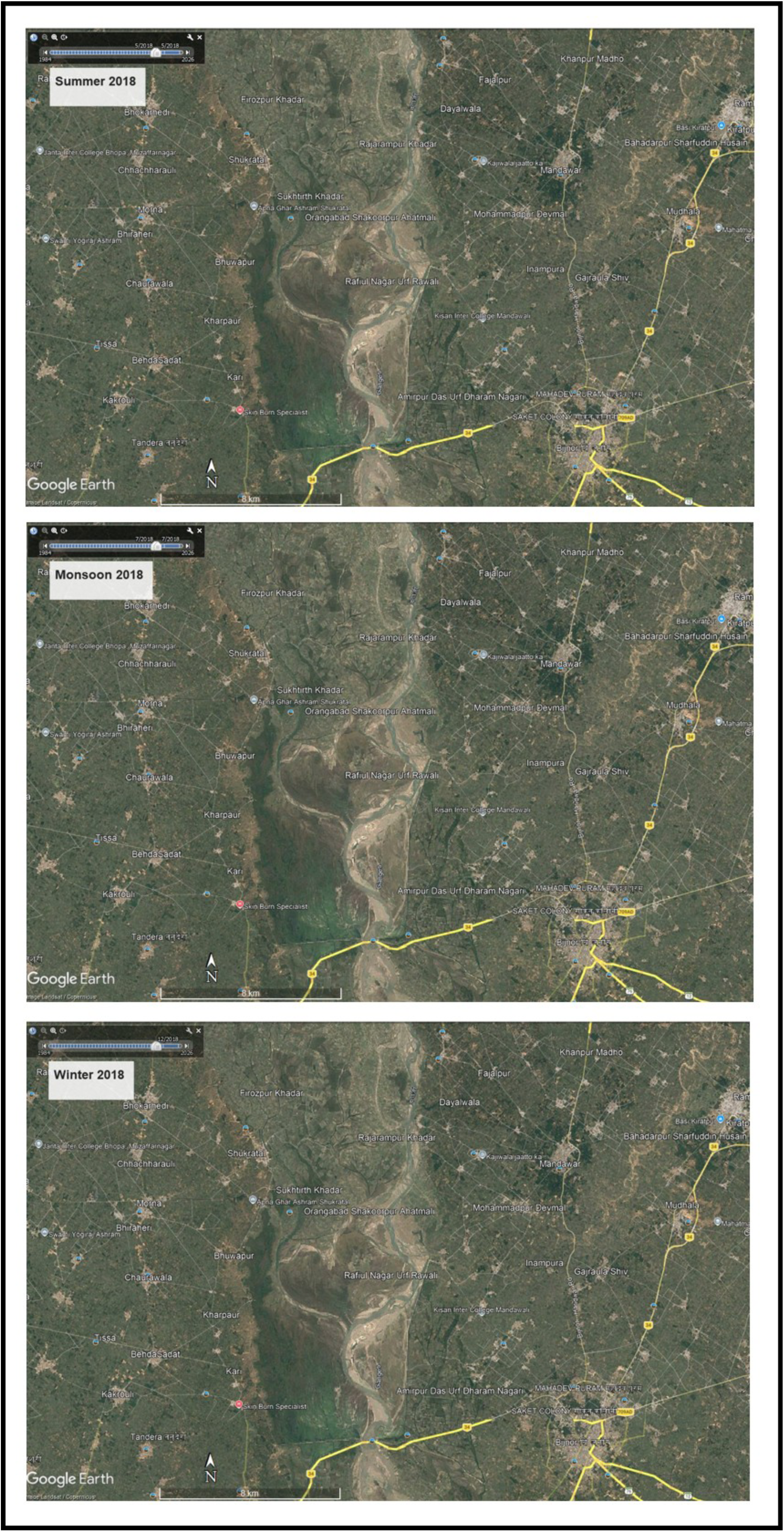
Google Earth Pro imagery of a representative grassland patch in the study area for the summer, monsoon, and winter seasons of 2018.

---

**Supplementary Table S1.** Details of Landsat images used for NDVI-based classification of grasslands.

| Satellite | No. of Tiles used | Path/Row | No. of Bands | Time Period | Grid cell size (m) |
| --- | --- | --- | --- | --- | --- |
| Landsat 8 OLI-TIRS | 2 | 146/39,146/40 | 11 | May 2018 | 30 |

**Supplementary Table S2.** Details of the accuracy assessment of four different vegetation type classifications.

| Vegetation types | Test plots | Correctly classified plots | Accuracy |
| --- | --- | --- | --- |
| Pure Saccharum (PS) | 27 | 26 | 96.20% |
| Saccharum Dominated Mix (SDM) | 49 | 41 | 84% |
| Phragmites-Typha Dominated Mix (PTDM) | 17 | 14 | 82.35% |
| Pure Phragmites-Typha (PPT) | 18 | 15 | 83.30% |
|  | 111 | 96 | **86.40%** |

## Notes

### Competing Interest Statement

The authors have declared no competing interest.

